# Multifunctional Rare-Earth-Ion-Doped Si-HAp Platforms Modulate Human BMSC Lineage-Associated Molecular Responses Without Enhancing Terminal Differentiation

**DOI:** 10.64898/2026.08.25.746707

**Authors:** Ariadna Pielok, Klaudia A. Marcinkowska, Natalia Charczuk, Joanna Sulecka-Zadka, Rafał J. Wiglusz, Agnieszka Śmieszek

## Abstract

**Introduction:** Advanced biomaterials for regenerative medicine are increasingly expected to combine multifunctionality and compatibility with tissue-specific cellular processes. In this context, hydroxyapatite-based platforms modified through ionic substitution represent promising candidates, as they may integrate structural similarity to bone mineral with additional biological functionality and luminescent properties, enabling diagnostic applications and real-time monitoring.

In this study, we evaluated whether silicate-phosphate substituted calcium hydroxyapatite Ca_10_(PO_4_)6-x(SiO_4_)x(OH)_2_ (where x = 1.5) co-doped with lithium(I), europium(III), and gadolinium(III) ions (Si-HAp-LEG) affects the osteogenic, chondrogenic, and adipogenic differentiation potential of human bone marrow stromal/stem cells (BMSCs).

**Methods:** Human BMSCs were cultured under lineage-specific differentiation conditions in the presence of undoped silicate-substituted phosphate hydroxyapatite (abbr. as Si-HAp), which served as a control, and two distinct Si-HAp-LEG formulations differing in gadolinium(III) (Gd^3+^) as well as lithium (Li^+^) and europium(III) (Eu^3+^) ion concentrations: Si-HAp-LEG-221 (1 mol% Gd^3+^ ion) and Si-HAp-LEG-222 (2 mol% Gd^3+^ ion). Differentiation-associated phenotypic outcomes, including extracellular matrix formation and lipid accumulation, were evaluated using Safranin O, Alizarin Red, and Oil Red O staining. In parallel, biomaterial-induced molecular responses were characterized at the transcriptomic and protein levels using RT-qPCR for selected coding and non-coding RNAs and Western blot analysis for representative lineage-associated proteins.

**Results:** Histochemical evaluation confirmed that, across all tested biomaterial groups, BMSCs retained the ability to form mineralized calcium deposits, proteoglycan-rich extracellular matrix, and intracellular lipid accumulation under osteogenic, chondrogenic, and adipogenic conditions, respectively. Quantitative staining analysis revealed no significant Si-HAp-LEG-dependent enhancement of terminal differentiation outcomes compared with undoped Si-HAp. In turn, the molecular response differed between biomaterials’ modifications. Si-HAp-LEG-222 induced the most prominent changes in transcriptional and post-transcriptional regulators, particularly within BMP/SMAD-associated pathways under osteogenic and chondrogenic conditions, underlying a potential link between gadolinium concentration and osteogenic lineage commitment. However, these transcriptomic responses were not mirrored by consistent changes at the protein level. The results suggest that silicate-phosphate substituted hydroxyapatite co-doped with Li+, Eu3+, and Gd3+ ions primarily affects the early regulatory pathways associated with BMSCs differentiation rather than enhancing their terminal maturation.

**Discussion:** In conclusion, the collective data indicate that Li^+^, Eu^3+^, and Gd^3+^ ions LEG co-doping broadens the multifunctional potential of Si-HAp by introducing imaging-related properties while preserving its underlying pro-regenerative character. Li^+^, Eu^3+^, and Gd^3+^ ions co-doped LEG-substituted Si-HAp may therefore be considered a compatible biomaterial platform that maintains BMSC cellular plasticity and supports balanced, differentiation-dependent modulation of lineage-associated molecular responses.

## 1 Introduction

The growing complexity of regenerative medicine has increased the demand for versatile biomaterial platforms capable of supporting different tissue-specific cellular responses while simultaneously providing additional diagnostic or theranostics functionality (Burkett et al., 2023; Keshari et al., 2026). Rather than acting solely as passive structural substitutes, such materials should interact with stem and progenitor cells, preserve their differentiation competence, and modulate lineage-associated signaling without disrupting the tightly coordinated processes that govern tissue repair (Vallet-Regí et al., 2026).

This is particularly relevant for multifunctional platforms intended for broad regenerative applications, where biological performance should be maintained across distinct mesenchymal lineages derived from multipotent cells such as bone marrow stromal/stem cells (BMSCs). Those cells contribute to bone and cartilage repair through osteogenic and chondrogenic differentiation, while their adipogenic potential reflects both the maintenance of multipotency and the balance between competing lineage programs within the bone marrow niche (Derubeis and Cancedda, 2004; An et al., 2023).

Accordingly, the design of advanced biocompatible scaffolds should extend beyond passive support of tissue regeneration and focus on actively directing BMSC fate toward therapeutically desirable and functional cells (Teng et al., 2026). In particular, materials that promote osteogenic and chondrogenic differentiation while limiting adipogenic commitment may offer a distinct advantage in strategies aimed at bone and cartilage regeneration (Fujimoto et al., 2022; Han et al., 2026; Vallet-Regí et al., 2026). Thus, the design of any biocompatible scaffold needs to be centered on improving the natural regenerative potential of the designated tissue and on patient safety.

Biomaterials based on synthetic nanosized hydroxyapatite (abbr. as nHAp) materials are intensively studied and utilized in various applications such as carriers for controlled drug delivery and bioactive scaffolds (Mo et al., 2023; Mondal et al., 2023) Nanosized hydroxyapatite is an extremely suitable material for bone tissue engineering due to its biocompatibility, exceptional mechanical, physical, and chemical properties, as well as osteoinductive activity and osseointegration (Abere et al., 2022). Furthermore, the hydroxyapatite crystal lattice readily accommodates ionic substitutions, allowing its physicochemical and biological properties to be tailored for specific applications (Ptáček, 2016). The co-doping of rare earth (RE^3+^) ions such as europium(III) (Eu^3+^) and gadolinium(III) (Gd^3+^) has been intensively studied due to their luminescent and magnetic properties, which warrant diagnostic possibilities and real-time monitoring of biomaterial incorporation and degradation at the implantation site(Namkhai and Jang, 2022; Pan et al., 2025). Furthermore, lithium ions (Li^+^) have also been reported to present beneficial bioactivity; similarly to Eu^3+^ and Gd^3+^ ions, they can promote osteogenesis and bone regeneration (Chen et al., 2024; Qi et al., 2025). Moreover, previous studies, together with our findings, indicate that lanthanide ions can enhance mesenchymal stem cell viability and exert anti-apoptotic effects, although these responses are strongly concentration-dependent (Charczuk et al., 2024; Zhu et al., 2019a; Liu et al., 2026; Feng et al., 2024; Wu et al., 2016; Cai et al., 2023; Zhao et al., 2019; Huang et al., 2021; Arioka et al., 2014; Geng et al., 2015; Lai et al., 2023).

In particular, our previous study focused on the biocompatibility and cytotoxicity of a multifunctional silicate-phosphate hydroxyapatite platform co-doped with Li⁺, Eu³⁺, and Gd³⁺ ions (abbr. as Si-HAp-LEG), which enabled the selection of Si-HAp-LEG-221 and Si-HAp-LEG-222 as the two most promising formulations (Charczuk et al., 2024). These platforms enhanced BMSC viability, exerted anti-apoptotic effects, modulated the expression of osteogenesis-related markers, including *RUNX2*, *TWIST1*, and *lnc-DANCR*, and displayed luminescent properties enabling *in vitro* tracking.

Building on these findings, the present study aimed to determine how Si-HAp-LEG-221 and Si-HAp- LEG-222 influence human BMSC lineage commitment toward osteogenic, chondrogenic, and adipogenic fates, with particular emphasis on the underlying regulatory mechanisms. This approach allowed us to assess whether these multifunctional platforms selectively direct BMSC differentiation toward therapeutically desirable lineages and to identify the most promising formulation for further preclinical development.

## 2 Materials and methods

### 2.1. Assessment of the Pro-Regenerative Properties of of Si_HAp-LEG platforms

#### 2.1.1. Biomaterials preparation and characterization

The undoped silicate-phosphate hydroxyapatite (abbr. as Si-HAp) as well as samples co-doped with Li^+^, Eu^3+^, and Gd^3+^ ions (abbr. as Si-HAp-LEG-221 and Si-HAp-LEG-222) were synthesized using the microwave-assisted hydrothermal method according to the procedure fully described in our previous work (Charczuk et al., 2024). The following substrates were used: Ca(NO_3_)_2_⋅4 H_2_O (Sigma-Aldrich), (NH_4_)_2_HPO_4_ (Acros Organics), TEOS (Alfa Aesar), Eu_2_O_3_ (Alfa Aesar), Gd_2_O_3_ (Alfa Aesar), and LiNO_3_ (Acros Organics). Briefly, rare-earth oxides (Eu_2_O_3_, Gd_2_O_3_) were digested in concentrated HNO_3_ (65%) and combined with the remaining precursors. The mixture was placed in a Teflon vessel. After adjusting the pH to ∼10 with NH_3_·H_2_O, the reaction was carried out at 240-250 °C under autogenous pressure (42-25 bar) for 90 minutes. The dried powders were calcined at 600°C for 3 hours.

#### 2.1.2. BMSC Culture in the Presence of Biomaterials under Osteogenic, Chondrogenic, and Adipogenic Differentiation Conditions

Human BMSCs were obtained from Merck Life Science (SCC034; Poznań, Poland), purchased at passage 2 (P2), and used to evaluate the pro-regenerative properties of the tested biomaterials. Cells in the logarithmic growth phase were detached, counted, and seeded into tissue culture plates at a density of 1.5 × 10⁴ cells/cm². After reaching 60–80% confluence, the tested biomaterials were introduced to the cultures, and trilineage differentiation toward the osteogenic, chondrogenic, and adipogenic lineages was initiated using commercially available StemPro^®^ Differentiation Kits (Thermo Fisher Scientific), according to the manufacturer’s instructions. BMSCs were maintained as adherent monolayer cultures in the respective complete StemPro^®^ Osteogenesis, Chondrogenesis, or Adipogenesis Differentiation Medium at 37°C in a humidified atmosphere containing 5% CO₂. Lineage-specific differentiation media were replaced every 2-4 days. Osteogenic and chondrogenic differentiation was continued for 20 days, whereas adipogenic differentiation was carried out for 10 days. At the end of the differentiation period, the cultures were subjected to lineage-specific staining and molecular analyses to comprehensively evaluate extracellular matrix deposition, lineage-specific phenotypic features, and molecular changes associated with BMSC differentiation.

#### 2.1.3. Histochemical Assessment of Biomaterial-Induced Lineage-Specific Extracellular Matrix Formation

Lineage-specific histochemical changes induced during osteogenic, chondrogenic, and adipogenic differentiation of BMSCs in the presence of the tested biomaterials were assessed using appropriate staining methods. Following the differentiation period, BMSC cultures were fixed with 4% paraformaldehyde (PFA) for 15 min at room temperature. Osteogenic differentiation was evaluated by Alizarin Red staining to detect mineralized calcium deposits, whereas chondrogenic differentiation was assessed using Safranin O staining to visualize proteoglycan-rich extracellular matrix. Adipogenic differentiation was evaluated by Oil Red O staining to detect intracellular lipid accumulation. The detailed staining procedures have been described previously (Zimoch-Korzycka et al., 2016; Śmieszek et al., 2018). The quantitative analysis of stain-positive areas in BMSC cultures incubated with the studied biomaterials was performed on the green channel (Color, Split Channels) using the Auto Threshold function (Yen thresholding algorithm) and Analyze Particles Function in ImageJ (ImageJ version 1.54p; National Institutes of Health, Bethesda, MD, USA). All reagents used for histochemical analyses were purchased from Merck (Poznań, Poland). Following staining, the cultures were examined using a Zeiss Primo Vert (Zeiss, Oberkochen, Germany) inverted microscope equipped with an Axiocam 208 color camera (Zeiss, Oberkochen, Germany), and representative images were acquired.

#### 2.1.4 Molecular Characterization of Biomaterial-Mediated Regulation of Lineage-Specific Differentiation Markers

To investigate how the tested biomaterials modulate the molecular programs associated with tissue-specific BMSC differentiation, the expression profiles of selected mRNA and non-coding RNA markers were analyzed following osteogenic, chondrogenic, and adipogenic induction in the presence of Si-HAp-LEG. Total RNA was isolated from human BMSC cultures subjected to the respective differentiation conditions using the phenol–chloroform extraction method, as originally described by Chomczynski et al. (Chomczynski and Sacchi, 1987). For this purpose, cultures were homogenized with TRI Reagent^®^ and processed according to the manufacturer’s instructions (Merck, Poznań, Poland).

The concentration and purity of the isolated RNA were assessed spectrophotometrically using Denovix (DS-11 Fx, Wilmington, DE, USA) based on absorbance measurements and the A260/A280 ratio. Genomic DNA was removed using the RNase-free PrecisionDNase kit (Primerdesign; BLIRT DNA, Gdańsk, Poland). Purified RNA was used for complementary DNA (cDNA) synthesis using the Tetro cDNA Synthesis Kit (Bioline Reagents Limited, London, UK) for mRNA analysis and the Mir-X™ miRNA First-Strand Synthesis Kit (Takara Bio Europe, Saint-Germain-en-Laye, France) for miRNA analysis. Genomic DNA digestion and reverse transcription were performed using a T100 Thermal Cycler (Bio-Rad, Hercules, CA, USA). The expression of selected mRNA and non-coding RNA markers associated with osteogenic, chondrogenic, and adipogenic differentiation was quantified using specific primer sets, the sequences of which are provided in Table S1. RT-qPCR analyses were performed using the CFX Connect™ Real-Time PCR Detection System (Bio-Rad, Hercules, CA, USA), with cycling conditions described previously (Smieszek et al., 2020). Expression data were normalized to glyceraldehyde-3-phosphate dehydrogenase (*GAPDH*) for mRNA targets and U6 for miRNA targets. Relative expression levels were calculated using the RQ_MAX_ algorithm and subsequently transformed to the log2 scale (Smieszek et al., 2020). To determine whether biomaterial-induced changes observed at the transcriptional level were accompanied by alterations in the abundance of selected proteins associated with osteogenic, chondrogenic, and adipogenic differentiation, Western blot analyses were performed as described previously (Smieszek et al., 2022). Briefly, BMSC cultures differentiated in the presence of biomaterials were lysed in ice-cold RIPA buffer supplemented with 1% protease and phosphatase inhibitor cocktail (Thermo Fisher Scientific, Warsaw, Poland). Total protein concentration was determined using the bicinchoninic acid assay (BCA; Thermo Fisher Scientific, Warsaw, Poland).

Protein samples were normalized to equal concentrations, mixed with 4× Laemmli sample buffer (Bio-Rad, Hercules, CA, USA), and heated at 95°C for 5 min. Equal amounts of protein were subsequently separated by 8–15% SDS-PAGE at 100 V for 90 min and transferred onto polyvinylidene difluoride (PVDF) membranes using 1× transfer buffer (Bio-Rad, Hercules, CA, USA). The membranes were blocked in 5% skim milk prepared in TBS-T and incubated with the respective primary antibodies overnight at 4°C. After washing with TBS-T, the membranes were incubated with appropriate secondary antibodies for 60 min at room temperature, then washed 5 times with TBS-T. Chemiluminescent signals were developed using DuoLuX^®^ Chemiluminescent and Fluorescent Peroxidase (HRP) substrate (Vector Laboratories; Biokom, Janki, Poland) and detected using the ChemiDoc XRS imaging system (Bio-Rad, Hercules, CA, USA). Densitometric analysis was performed using Image Lab™ software (version 6.1; Bio-Rad: Hercules, CA, USA; 2020) using an established protocol (Marcinkowska et al., 2026). Details of the primary and secondary antibodies used in the study are provided in Table S2.

### 2.2. Statistical analysis

Unless otherwise stated, experiments were performed using at least three independent biological replicates, with the number of technical replicates determined by the analytical method. Most analyses were performed in triplicate, whereas Western blot analyses were conducted in duplicate. Data distribution was assessed using the Shapiro–Wilk test, while homogeneity of variance was evaluated using Fisher’s test. Statistical differences among multiple groups were analysed using either one-way analysis of variance (ANOVA) or the Kruskal–Wallis test, depending on data distribution, followed by Tukey’s or Dunn’s post hoc multiple-comparison test, respectively. Statistical analyses and data visualisation were performed using GraphPad Prism 10 (GraphPad Software, La Jolla, CA, USA) and RStudio (ver. 2026.05.1). Data were presented as mean ± standard deviation (SD). Differences were considered statistically significant at p < 0.05 and denoted as follows: *p < 0.05, **p < 0.01, and ***p < 0.001.

## 3 Results

### 3.1 LEG-Doping Modulates Chondrogenesis-Related Molecular Signaling Without Enhancing Mature Matrix Formation

Histochemical assessment using Safranin O staining revealed the formation of proteoglycan-rich extracellular matrix in all BMSC cultures subjected to chondrogenic differentiation (Figure 1A). Quantitative analysis of Safranin O staining revealed no statistically significant differences among the analyzed groups (Figure 1B). Nevertheless, both Si-HAp-LEG-221 and Si-HAp-LEG-222 exhibited lower mean Safranin O-positive areas than undoped Si-HAp, indicating a non-significant trend toward reduced deposition of proteoglycan-rich extracellular matrix following ions co-doping.

**Figure 1.**
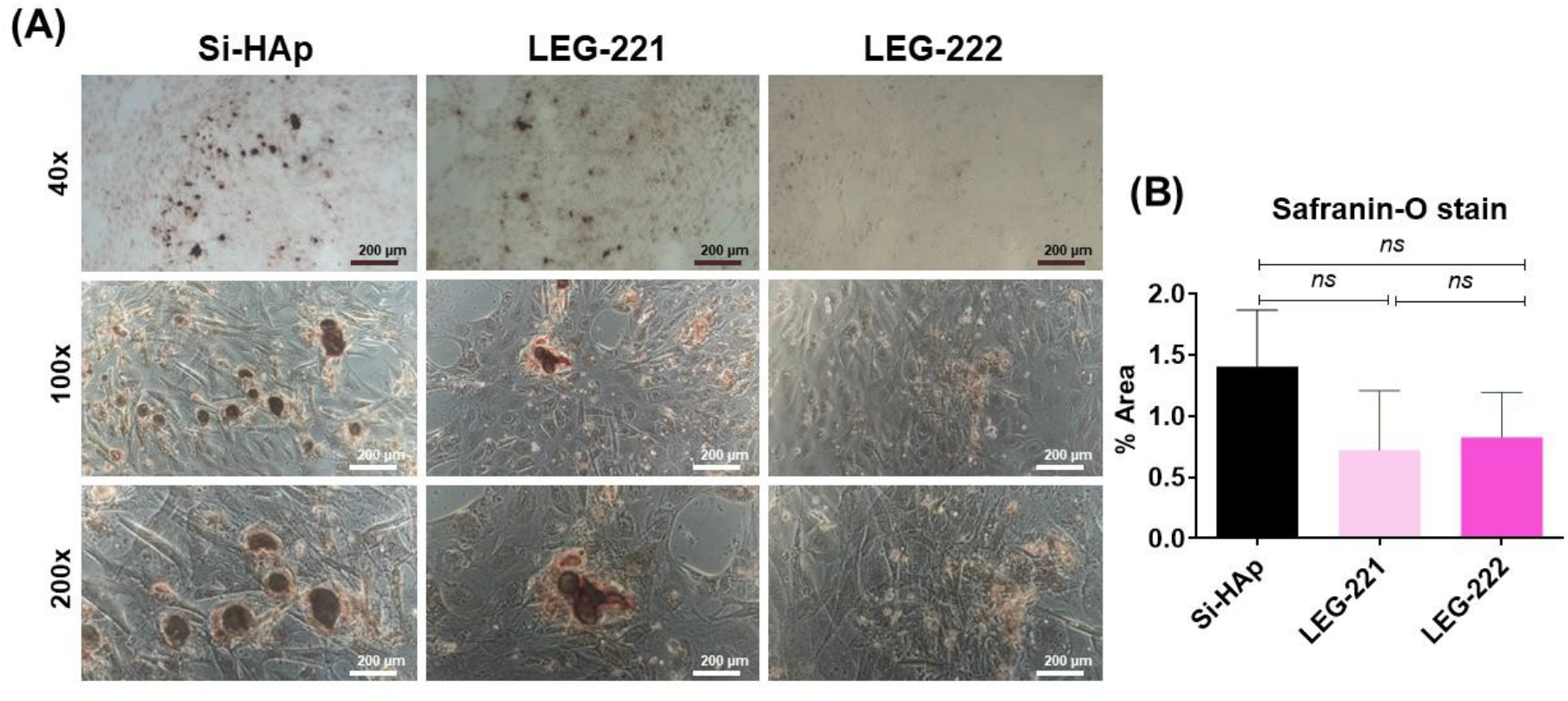
Effects of rare-earth-ion-doped Si-HAp on proteoglycan-rich extracellular matrix formation during chondrogenic differentiation of BMSCs. (A) Representative microphotographs of Safranin O-stained BMSC cultures maintained under chondrogenic conditions in the presence of the tested biomaterials, including the Si-HAp pure platform and matrices co-doped with Si-HAp-LEG-221 and Si-HAp-LEG-222. Images were acquired at 40×, 100×, and 200× magnification. Scale bars are indicated in the microphotographs. (B) Quantitative analysis of the Safranin O-positive area revealed no significant differences among the experimental groups. The analysis was performed using the ImageJ application based on a minimum of two microphotographs per technical replicate. Data were obtained from three independent biological replicates, each assessed in two technical replicates. Results are presented as columns with bars representing mean value ± SD. Non-statistical differences were indicated with “ns”.

Molecular profiling revealed a distinct response of BMSCs cultured under chondrogenic conditions in the presence of the tested biomaterials, with Si-HAp-LEG-222 eliciting the most pronounced changes (Figure 2). This formulation significantly increased the expression of multiple components of BMP/SMAD-associated signalling, including *BMPR1A*, *BMPR2*, *SMAD1*, *SMAD2*, *SMAD3*, and *SMAD4*, accompanied by increased expression of lineage-regulatory markers such as *RUNX2*, *TWIST1*, and *DANCR*. In contrast, the expression of several markers associated with cartilage extracellular matrix formation, including *ACAN* and *COL2*, remained unchanged in BMSC at all tested conditions.

**Figure 2.**
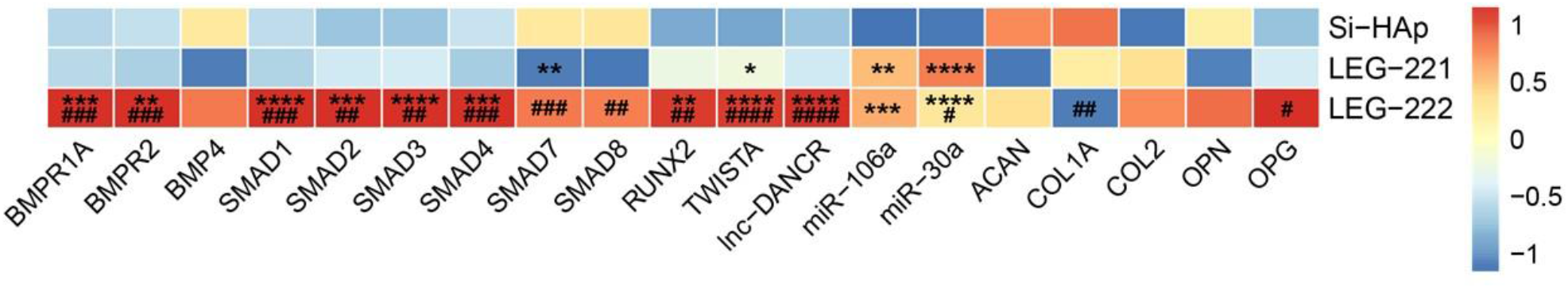
Data from RT-qPCR analysis of selected mRNAs, miRNAs, and lncRNAs in BMSCs cultured under chondrogenic conditions in the presence of studied platforms. Results are illustrated using a heatmap generated in RStudio (ver. 2026.05.1) using the pheatmap package, based on mean relative quantification (RQ) values on a log scale. The colour scale represents row-wise z-scores, ranging from −1 (dark blue; lower relative expression) to 1 (red; higher relative expression), with yellow indicating the mean expression level (z-score = 0). Statistical differences between the cultures incubated with Si-HAp and the cultures incubated with Si-HAp-LEG-221/Si-HAp-LEG-222 platforms are indicated with asterisks (*p < 0.05, **p < 0.01, ***p < 0.001, and ****p < 0.0001) while statistical differences between the BMSCs incubated with Si-HAp-LEG-221 and BMSCs incubated with Si-HAp-LEG-222 are indicated with hashtags (#p < 0.05, ##p < 0.01, ###p < 0.001, and ####p < 0.0001). Only statistically significant differences are marked on the heatmap; unmarked tiles represent comparisons where no statistically significant differences were observed.

This transcriptional response was accompanied by a distinct ions-co-doping-dependent miRNA expression profile, including changes in *miR-30a* and *miR-106a*, suggesting an additional layer of post-transcriptional regulation of lineage-associated processes.

Although Si-HAp-LEG-222 induced a coordinated molecular response involving BMP/SMAD signalling and lineage-regulatory pathways, this effect does not correspond with enhanced deposition of proteoglycan-rich extracellular matrix, as indicated by the Safranin O analysis.

The obtained findings suggest that Si-HAp co-doping with Li^+^, Eu^3+^, and Gd^3+^ ions, particularly in the Si-HAp-LEG-222 formulation, exerts a pronounced effect on BMP/SMAD-associated signalling and early lineage-regulatory programs.

However, this molecular response is not accompanied by enhanced expression of mature cartilage matrix markers or increased deposition of proteoglycan-rich extracellular matrix. This interpretation is further supported by Western blot analysis, which revealed no significant ions co-doping-dependent alterations in the accumulation of ACAN (Figure 3C), OPN protein (Figure 3D), COL1 (Figure 3E), OPG (Figure 3F), and COL2 (Figure 3G). Moreover, the intracellular expression of RUNX2 protein (Figure 3B) was not altered by biomaterial exposure, regardless of whether the cells were cultured with Si-HAp, Si-HAp-LEG-221, or Si-HAp-LEG-222 (Figure 3 B).

**Figure 3.**
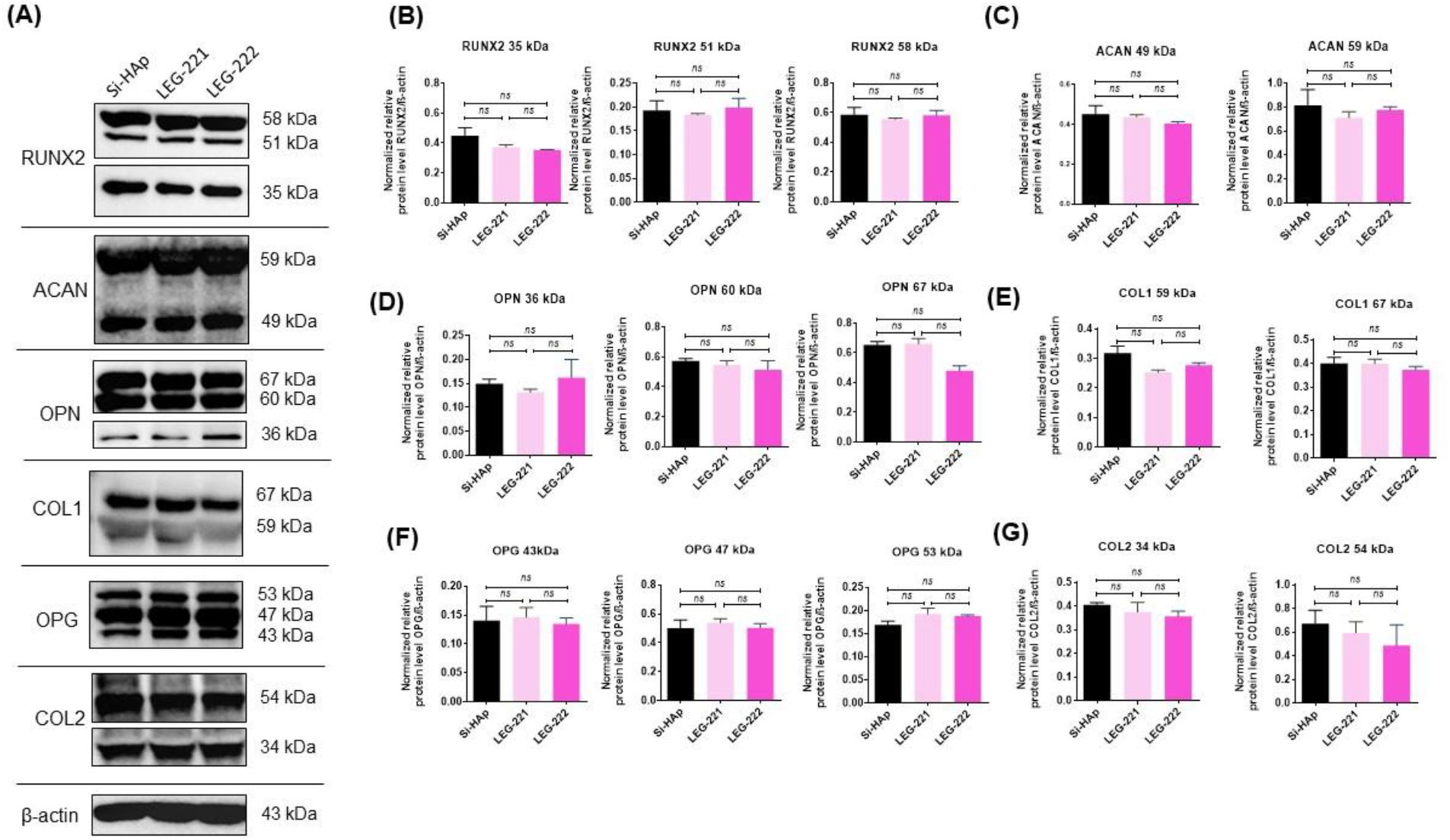
The results of immunodetection of intracellularly accumulated proteins. The representative immunoblots (A) show bands characteristic of detected proteins. A comparative analysis was performed to assess differences in the expression of markers associated with lineage regulation and late chondrogenesis, including RUNX2 (B), ACAN (C), OPN (D), COL1 (E), OPG (F), and COL1 (G). Results are presented as columns with bars representing mean value ± SD. Non-significant statistical differences are indicated with the “ns” symbol.

### 3.2 LEG Co-Doping Influences Early Osteogenic Regulatory Networks Without Enhancing Terminal Matrix Mineralization

Alizarin Red staining revealed the formation of mineralized osteogenic nodules in BMSCs cultured under osteogenic conditions in the presence of all tested biomaterials (Figure 4 A4A). Quantitative analysis showed a comparable degree of matrix mineralization among the groups, with no statistically significant differences detected between undoped Si-HAp, Si-HAp-LEG-221, and Si-HAp-LEG-222 (Figure 4B). Although the Si-HAp-LEG-222 group displayed a numerically greater Alizarin Red-positive area, this difference did not reach statistical significance. Thus, LEG the ions co-doping did not result in a measurable enhancement of mineralized extracellular matrix deposition under the applied experimental conditions.

**Figure 4.**
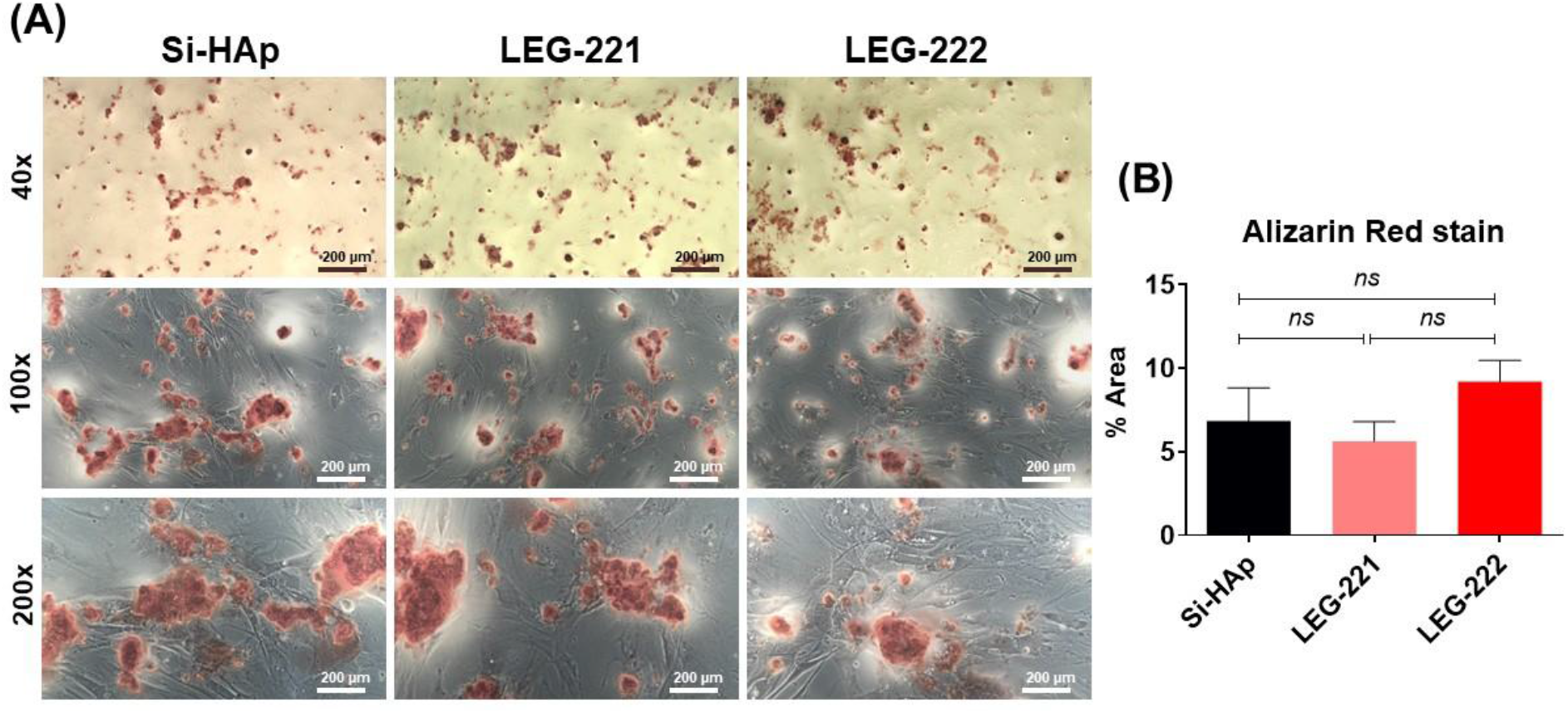
Effects of rare-earth-ion-co-doped Si-HAp on calcium-rich extracellular matrix formation during osteogenic differentiation of BMSCs. **(A)** Representative microphotographs of Alizarin Red-stained BMSC cultures maintained under osteogenic conditions in the presence of the tested biomaterials, including an undoped Si-HAp platform and matrices co-doped with Si-HAp-LEG-221 and Si-HAp-LEG-222. Images were acquired at 40×, 100×, and 200× magnification. Scale bars are indicated in the microphotographs. **(B)** Quantitative analysis of the Alizarin Red-positive area revealed no significant differences among the experimental groups. The analysis was performed using the ImageJ application based on a minimum of two microphotographs per technical replicate. Data were obtained from three independent biological replicates, each assessed in two technical replicates. Results are presented as columns with bars representing mean value ± SD. Non-statistical differences were indicated with the “ns”.

Despite the absence of a clear functional effect on matrix mineralization, molecular profiling revealed substantial ions-codoping-dependent differences in the expression of osteogenesis-associated regulatory markers (Figure 5). The most pronounced transcriptional response was observed in the Si-HAp-LEG-222 group and involved multiple components of BMP/SMAD-associated signalling and lineage-regulatory programs, including *BMPR1A*, *BMPR2*, *SMAD* family members, *ALPL*, *RUNX2*, *BMP4*, *COL1A1*, and *DANCR* (Figure 5). In contrast, Si-HAp-LEG-221 generally induced a more limited and less coordinated transcriptional response. The analysed miRNA profile further distinguished the experimental groups, with Si-HAp-LEG-222 showing a marked reduction in *miR-106a* expression and lower *miR-30a* levels, indicating an additional post-transcriptional component of the biomaterial-induced response (Figure 5).

**Figure 5.**
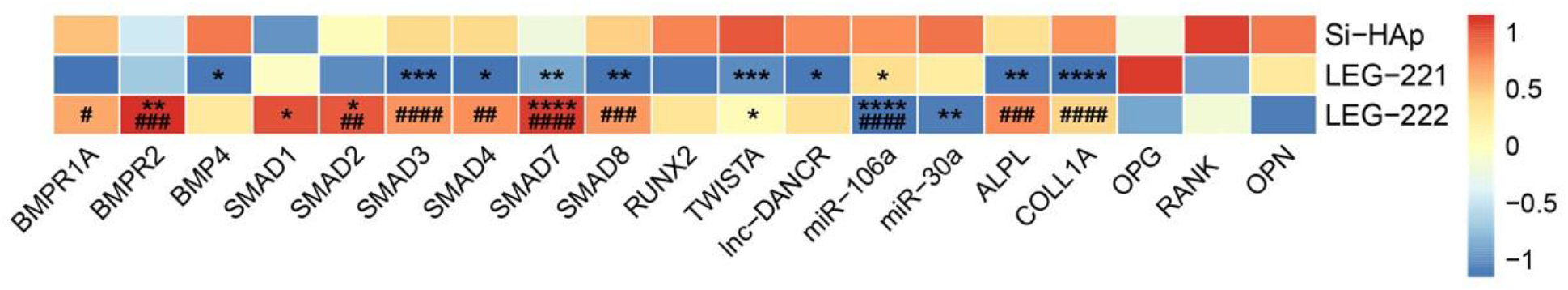
Data from RT-qPCR analysis of selected mRNAs, miRNAs, and lncRNAs in BMSCs cultured under osteogenic conditions in the presence of studied platforms. Results are illustrated using a heatmap generated in RStudio (ver. 2026.05.1) using the pheatmap package, based on mean relative quantification (RQ) values on a log scale. The colour scale represents row-wise z-scores, ranging from −1 (dark blue; lower relative expression) to 1 (red; higher relative expression), with yellow indicating the mean expression level (z-score = 0). Statistical differences between the cultures incubated with Si-HAp and the cultures incubated with Si-HAp-LEG-221/Si-HAp-LEG-222 platforms are indicated with asterisks (*p < 0.05, **p < 0.01, ***p < 0.001, and ****p < 0.0001) while statistical differences between the BMSCs incubated with Si-HAp-LEG-221 and BMSCs incubated with Si-HAp-LEG-222 are indicated with hashtags (#p < 0.05, ##p < 0.01, ###p < 0.001, and ####p < 0.0001). Only statistically significant differences are marked on the heatmap; unmarked tiles represent comparisons where no statistically significant differences were observed.

Importantly, the transcriptional changes observed in cultures on the ions co-doped platforms were not accompanied by corresponding alterations in the accumulation of major osteogenesis-associated proteins (Figure 6A). Western blot analysis revealed no significant the ions co-doping-dependent differences in BMP2 (Figure 6B), COL1(Figure 6D), OPN Figure 6E), or OPG protein levels (Figure 6F). RUNX2 protein accumulation also remained largely unchanged, except for a significant reduction in the 34-kDa band in the Si-HAp-LEG-221 group, whereas the 53- and 60-kDa bands showed no significant differences among the tested conditions (Figure 6C).

**Figure 6.**
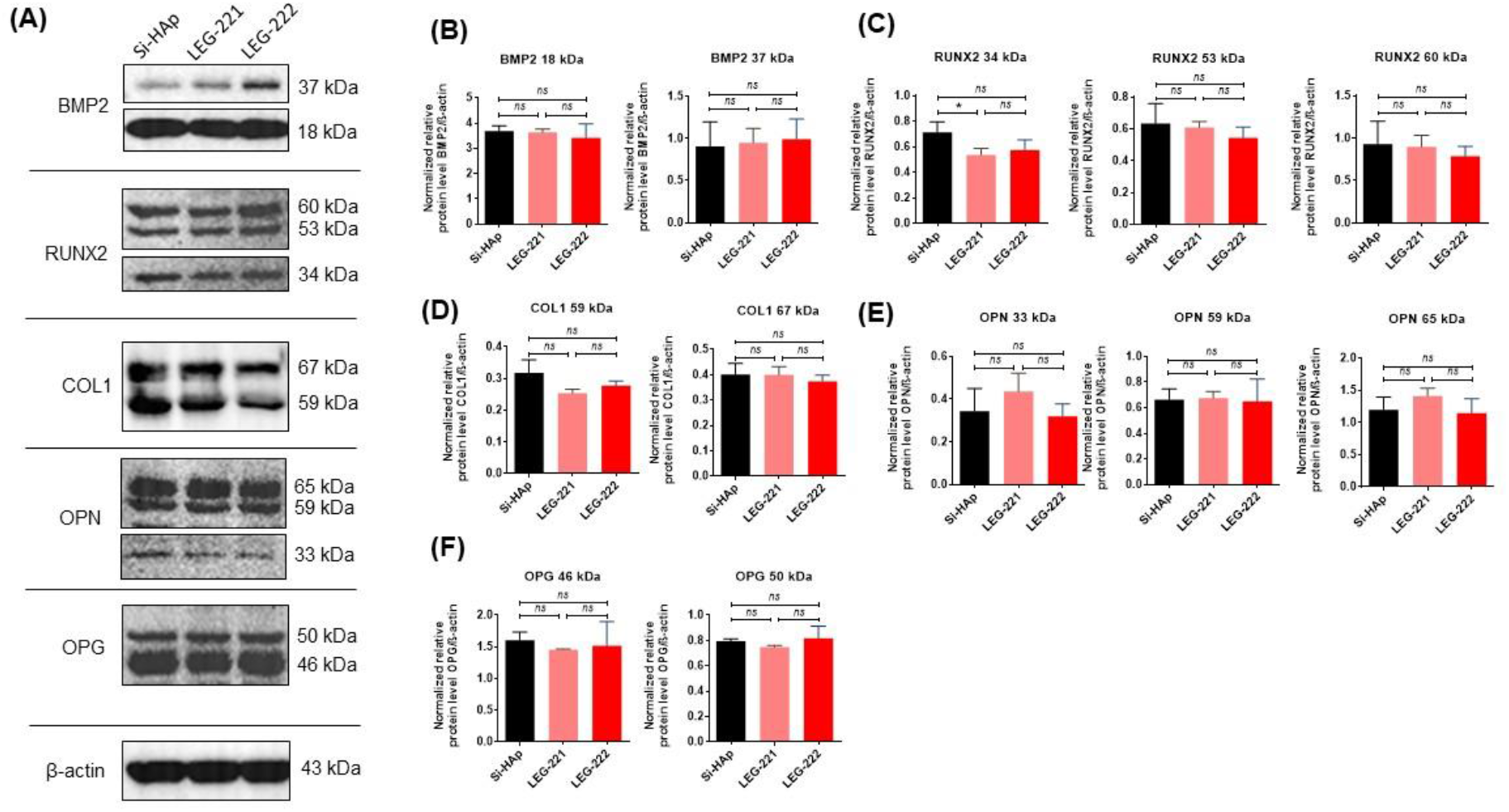
The results of the immunodetection of intracellular proteins under osteogenic conditions. Representative immunoblots (A) show bands corresponding to the detected proteins. A comparative analysis was performed to assess differences in the expression of markers associated with lineage regulation, extracellular matrix formation and maturation, including BMP2 (B), RUNX2 (C), COL1 (D), OPN (E) and OPG (F). Results are presented as columns with bars representing mean value ± SD. Statistical differences were indicated with an asterisk (* p-value < 0.05), while the “ns” symbol refers to a non-significant difference.

Taken together, these findings indicate that the ions co-doping, particularly in the Si-HAp-LEG-222 formulation, primarily reshapes osteogenesis-associated transcriptional and post-transcriptional programs without producing a corresponding increase in mature osteogenic protein accumulation or matrix mineralization.

### 3.3 The Ions Co-Doping Alters Adipogenesis-Associated Regulatory Profiles Without Enhancing Lipid Accumulation

Oil Red staining revealed intracellular lipid accumulation in BMSCs subjected to adipogenic differentiation in the presence of all tested biomaterials (Figure 7A). Although no statistically significant differences were detected among the groups, both LEG-doped formulations co-doping materials showed a numerically lower lipid-positive area than undoped siSi-HAp, indicating a non-significant trend toward reduced lipid accumulation.

**Figure 7.**
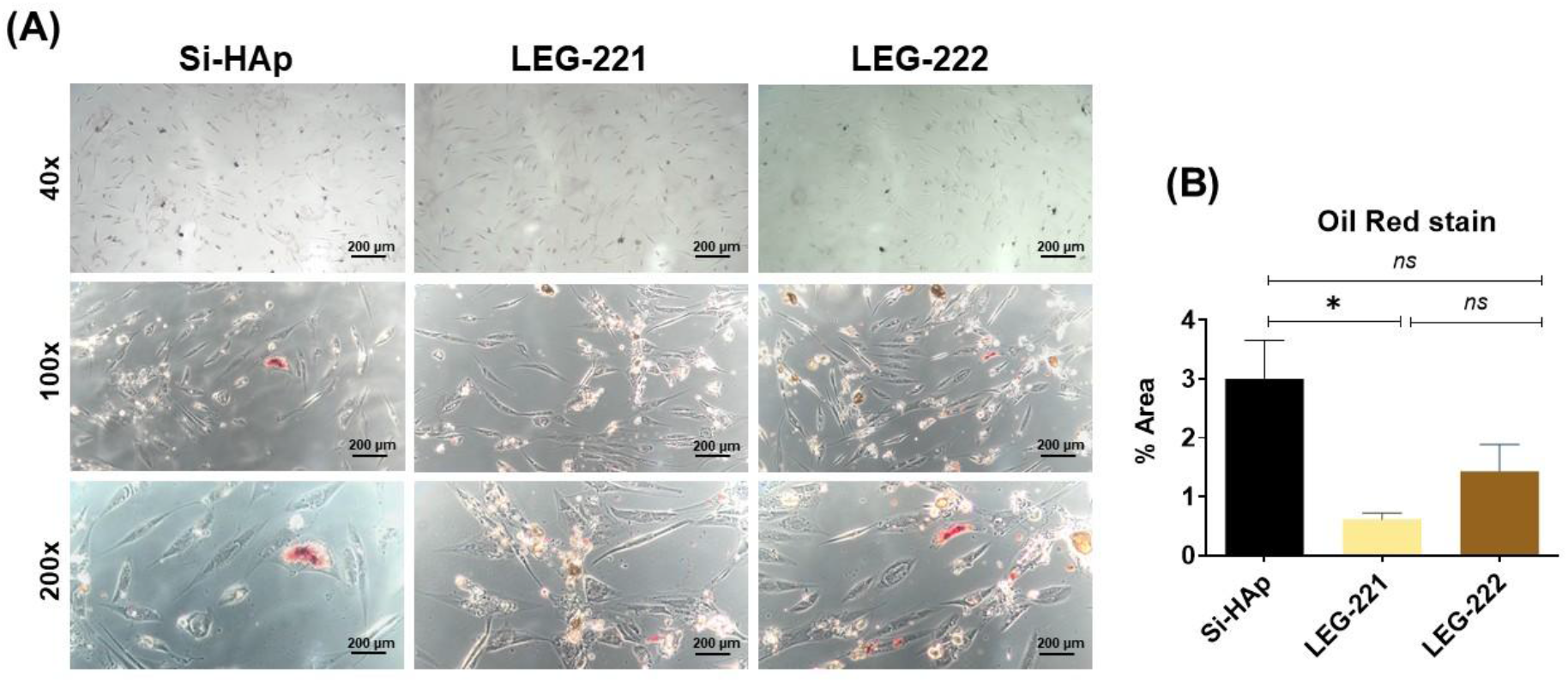
Effects of rare-earth-ion-co-doped Si-HAp on lipid accumulation during adipogenic differentiation of BMSCs. **(A)** Representative microphotographs of Oil Red-stained BMSC cultures maintained under osteogenic conditions in the presence of the tested biomaterials, including the undoped Si-HAp platform and matrices co-doped with the ions (Si-HAp-LEG-221 and Si-HAp -LEG-222). Images were acquired at 40×, 100×, and 200× magnification. Scale bars are indicated in the microphotographs. **(B)** Quantitative analysis of the Oil Red-positive area revealed no significant differences among the experimental groups. The analysis was performed using ImageJ applications based on a minimum of two microphotographs per technical replicate. Data were obtained from three independent biological replicates, each assessed in two technical replicates. Results are presented as columns with bars representing mean value ± SD. Statistical differences were indicated with an asterisk (* p-value < 0.05), while the “ns” symbol refers to a non-significant difference.

Gene-expression profiling revealed selective LEG-dependent changes in regulators associated with lineage commitment and adipogenesis (Figure 8). The most pronounced alterations were observed in the Si-HAp-LEG-222 group, which showed reduced *adiponectin* and *DANCR* expression, accompanied by a distinct shift in the expression of *TWIST1*. While *TWIST1* was markedly increased in the Si-HAp-LEG-221 group and reduced in the Si-HAp-LEG-222 group. In contrast, mRNA expression of *PPARγ*, *leptin*, *ACAN*, and *RUNX2* remained largely unchanged across the tested conditions.

**Figure 8.**
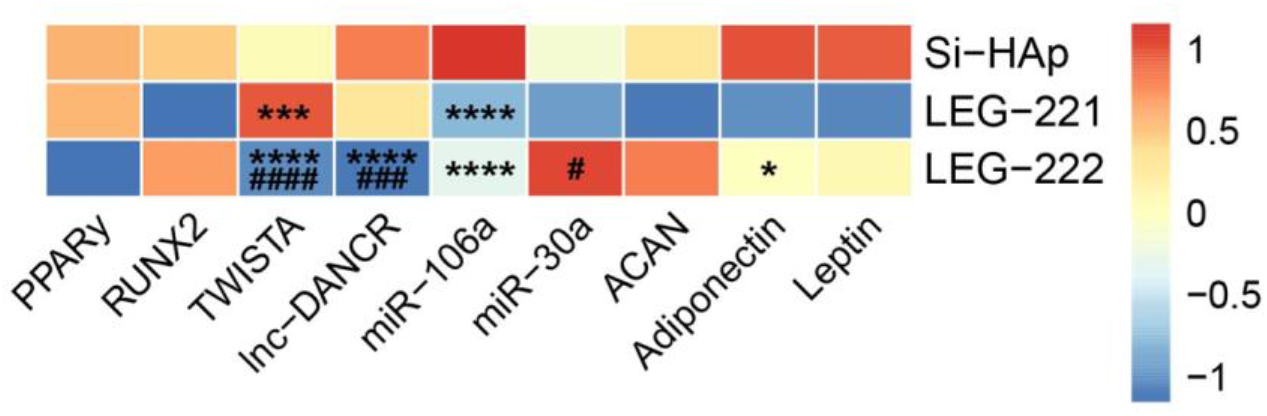
Data from RT-qPCR analysis of selected mRNAs, miRNAs, and lncRNAs in BMSCs cultured under adipogenic conditions in the presence of studied platforms. Results are illustrated using a heatmap generated in RStudio (ver. 2026.05.1) using the pheatmap package, based on mean relative quantification (RQ) values on a log scale. The color scale represents row-wise z-scores, ranging from −1 (dark blue; lower relative expression) to 1 (red; higher relative expression), with yellow indicating the mean expression level (z-score = 0). Statistical differences between the cultures incubated with Si-HAp and the cultures incubated with LEG-221/LEG-222 platforms are indicated with asterisks (*p < 0.05, **p < 0.01, ***p < 0.001, and ****p < 0.0001) while statistical differences between the BMSCs incubated with LEG-221 and BMSCs incubated with LEG-222 are indicated with hashtags (#p < 0.05, ##p < 0.01, ###p < 0.001, and ####p < 0.0001). Only statistically significant differences are marked on the heatmap; unmarked tiles represent comparisons where no statistically significant differences were observed.

The miRNA profile provided an additional layer of evidence for the ions co-doping-dependent molecular regulation. Both co-doped formulations markedly reduced *miR-106a* expression compared with undoped Si-HAp, whereas *miR-30a* displayed a formulation-dependent pattern, with higher expression in the Si-HAp-LEG-222 than in the Si-HAp-LEG-221 group. These findings indicate that the ions co-doping modifies not only transcriptional regulators but also post-transcriptional mechanisms potentially involved in BMSC lineage-associated responses.

Importantly, the observed transcriptional alterations were not accompanied by corresponding changes in the protein levels of adiponectin and leptin (Figure 9A). Western blot analysis revealed no significant differences in adiponectin (Figure 9C) or leptin (Figure 9D) accumulation among the tested groups, further supporting the absence of a clear LEG-dependent enhancement of the mature adipogenic phenotype. In addition, during adipogenic differentiation, the expression of the 36- and 62-kDa OPN bands was increased in the ions co-doping-treated groups. The highest levels were observed in the Si-HAp-LEG-222 group and were significantly higher than those in both the Si-HAp and Si-HAp-LEG-221 groups, whereas no significant difference was found between Si-HAp-LEG-221 and Si-HAp. In contrast, the expression of the 66-kDa OPN band did not differ significantly among the experimental groups.

**Figure 9.**
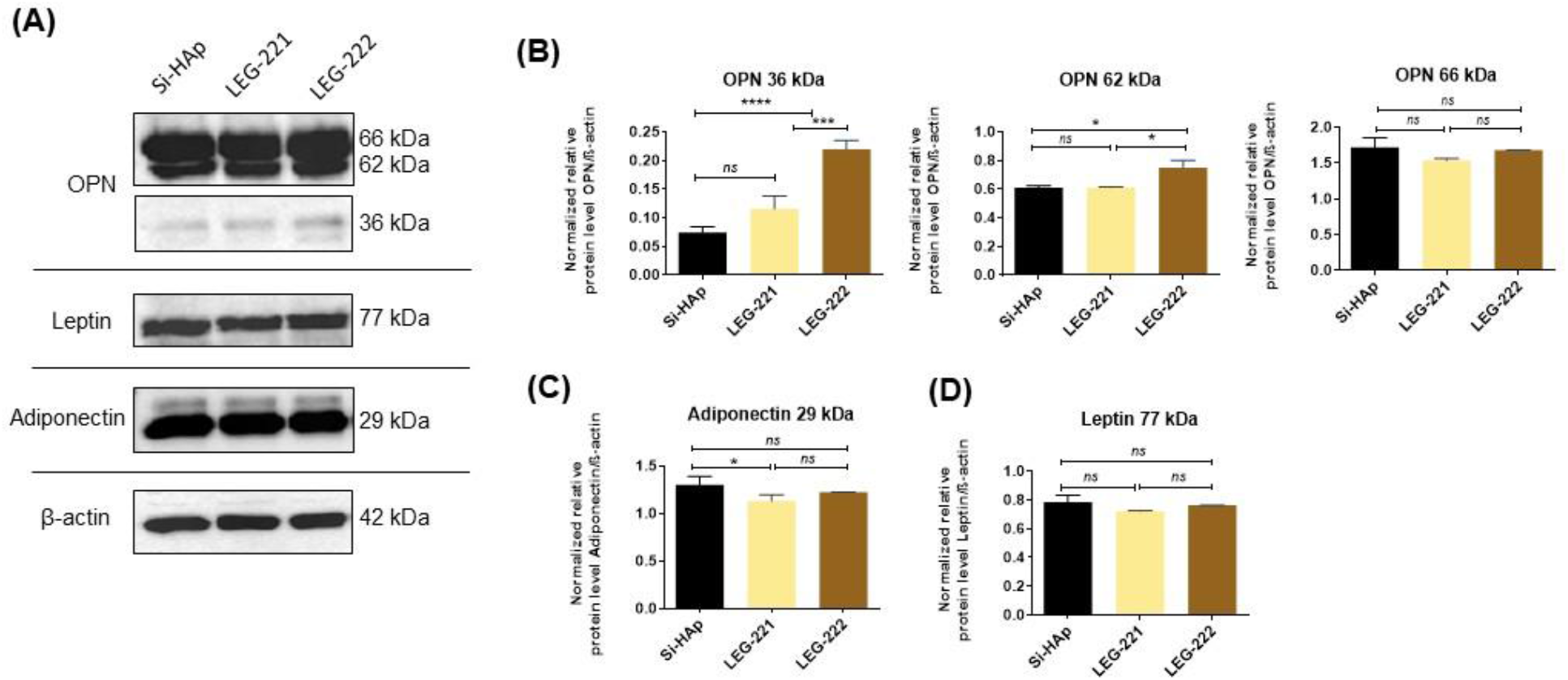
The results of immunodetection of intracellularly accumulated proteins in BMSCs cultured in adipogenic conditions. The representative immunoblots (A) show bands characteristic of detected proteins. A comparative analysis was performed to assess differences in the expression of markers associated with lineage commitment and adipocyte maturation, including OPN (B), adiponectin (C), and leptin (D). Results are presented as columns with bars representing mean value ± SD. Statistical differences were indicated with asterisks (* p-value < 0.05, *** p-value < 0.001, and ****p-value < 0.0001), while the “ns” symbol refers to a non-significant difference.

## 4 Discussion

A major challenge in the development of advanced biomaterials for regenerative medicine is to introduce additional diagnostic functionalities without compromising the intrinsic biological properties that make these materials suitable for tissue repair (Wang et al., 2026). Hydroxyapatite-based platforms are particularly attractive in this context because of their well-established biocompatibility and ability to support bone-related cellular responses (Mo et al., 2023; Mondal et al., 2023; Vallet-Regí et al., 2026). However, ion incorporation may alter cell–material interactions and consequently influence stem cell fate (Kersey et al., 2024). As we showed previously, Si-HAp co-doped with Li⁺, Eu³⁺, and Gd³⁺ ions preserved the platform’s cytocompatibility, allowing BMSCs to maintain proper morphology, cytoskeletal organization, and metabolic activity without inducing a pro-apoptotic transcriptional response. At the same time, the ion-doped materials were biologically active, as evidenced by their ability to modulate key regulators associated with lineage commitment, including *RUNX2*, *TWIST1*, and lncRNA *DANCR* (Charczuk et al., 2024).

Substituting co-dopants into the HAp crystal lattice introduces distinct multifunctional properties, including optical and magnetic functionalities relevant for bioimaging applications(Kersey et al., 2024). However, it also raises the important question of whether these added functions can be achieved without compromising the material’s ability to support lineage-specific BMSC differentiation.

Indeed, our findings indicate that the ions substitution, particularly in the Si-HAp-LEG-222, introduces pronounced changes in lineage-associated molecular regulation without substantially disturbing the terminal differentiation profile of BMSCs or compromising mature matrix formation. Importantly, we show that the additional multifunctionality conferred by Li⁺, Eu³⁺, and Gd³⁺ ions co-doping can be achieved while largely preserving the intrinsic biological performance of the Si-HAp platform. Rather than acting as a strong inducer of terminal differentiation, the Li⁺, Eu³⁺, and Gd³⁺ ions co-doped Si-HAp platform appears to selectively reshape lineage-associated regulatory programs, with Si-HAp-LEG-222 exerting the most pronounced molecular effects, reflected in coordinated, differentiation-dependent changes in the BMP/SMAD–RUNX2 axis and associated lncRNA and miRNA networks.

From the perspective of multifunctional biomaterial design, the phenotypic stability of BMSCs differentiated in the presence of pro-regenerative stimuli supports the use of the Li⁺, Eu³⁺, and Gd³⁺ ions co-doped Si-HAp in diverse regenerative settings. The Li⁺, Eu³⁺, and Gd³⁺ co-doping did not compromise BMSCs’ ability to undergo tissue-specific differentiation or to form lineage-appropriate mature structures.

Previous studies by Alicka et al. demonstrated that Li⁺ ions as well as Li⁺ ion-doped nHAp can enhance osteogenesis in human adipose-derived stromal/stem cells, as reflected by increased calcium deposition, modulation of GSK3β/β-catenin signalling, and altered expression of osteogenesis-associated markers, including BMP2, ALP, OCN, and OPN (Alicka et al., 2019). Importantly, the ion-co-doped formulations in the previous study were evaluated primarily with respect to luminescence and short-term cytocompatibility rather than their capacity to support terminal lineage differentiation. The present study extends these observations by demonstrating that Li⁺/Eu³⁺/Gd³⁺ ion co-doping not only preserves the low-cytotoxicity profile of the Si-HAp platform but, more importantly, does not affect the ability of human BMSCs to undergo tissue-specific osteogenic, chondrogenic, and adipogenic differentiation. The Li⁺, Eu³⁺, and Gd³⁺ ions co-doping appears to provide a fine-tuning effect on lineage-regulatory programs while preserving the regenerative competence of the undoped Si-HAp platform.

To our knowledge, this is the first study to comprehensively evaluate the effects of Li⁺/rare-earth-ion-doped nHAp on BMSC fate across three canonical mesenchymal differentiation pathways: osteogenesis, chondrogenesis, and adipogenesis.

In our study, we assessed a broad panel of markers associated with lineage-specific differentiation. However, in the context of nHAp functionalization, particular attention should be given to osteopontin (OPN), which was also highlighted by Alicka et al. as an important indicator of the biological response to Li⁺-containing nHAp (Alicka et al., 2019). As a multifunctional matricellular protein involved in cell–matrix interactions, mineralization, and tissue remodelling, OPN contributes to matrix regulation in both bone and cartilage. In the study by Alicka et al., Li⁺ ions, nHAp, and Li⁺-doped nHAp biomaterials differentially affected intracellular, extracellular, full-length, and MMP-cleaved OPN, with the highest extracellular level observed after exposure to free lithium ions.

In contrast, the present study revealed no significant the ions co-doping-dependent changes in OPN protein accumulation under either osteogenic or chondrogenic conditions. This stability, together with preserved mineralization and lineage-specific matrix formation, indicates that LEG-doping does not significantly affect OPN expression or matrix-remodeling components during BMSCs’ osteogenic and chondrogenic differentiation. The most distinct changes in OPN protein accumulation have been observed in BMSCs subjected to adipogenic differentiation in the presence of LEG-222. Notably, the changes in OPN protein accumulation in response to Si-HAp-LEG-222 were not accompanied by corresponding shifts in other crucial adipogenesis-related regulators or lineage-specific lipid droplet deposition.

An additional mechanistic perspective partially supporting our results was provided by Wang et al., who demonstrated that luminescent hydroxyapatite is a metabolically active regulator of osteogenic differentiation. In their model, defect-related luminescent HAp was internalized by BMSCs and localized within lysosomes, where its degradation promoted phosphate release. The resulting increase in intracellular ATP and extracellular adenosine was linked to activation of A2B receptor-dependent cAMP/PKA signalling, leading to increased expression of *RUNX2*, *BMP2*, *OCN*, and *COL1*, enhanced ALP activity, and increased matrix mineralization (Wang et al., 2016). These findings indicate that changes in the physicochemical properties of the HAp, even without incorporation of bioactive ions, may engage BMP-associated osteogenic transcriptional programs through upstream metabolic signalling.

BMP signalling, however, is not restricted to bone formation - it represents a broader regulatory network involved in tissue morphogenesis, repair, and lineage specification of mesenchymal progenitor cells (Kwon et al., 2013; Cogo et al., 2025; Jagdish et al., 2025). A conceptually related pattern was observed in our study, in which Si-HAp-LEG-222 induced coordinated transcriptional changes in *BMP* receptors and downstream *SMAD* regulators under osteogenic and chondrogenic conditions. Although these changes did not translate into enhanced terminal matrix formation, they suggest that incorporation of LEG-222 may influence the molecular competence of BMSCs to respond to lineage-specific differentiation cues.

The observed BMP/SMAD-associated transcriptional response was paralleled by a lineage-dependent profile of lncRNA *DANCR*. This long-noncoding RNA has previously been described as a potent regulator of osteogenic commitment through pathways converging on *RUNX2*, *Wnt*, and *p38 MAPK* signalling (Zhang et al., 2019; Liu et al., 2022). In our study, Si-HAp-LEG-222 induced a higher accumulation of this transcript in BMSC under osteogenic and chondrogenic conditions, but with a pronounced reduction during adipogenic differentiation. This pattern further supports the concept that the ions co-doped biomaterials differentially reshape lineage-regulatory programs.

The expression profile of *TWIST1*, a highly conserved transcription factor controlling mesenchymal lineage commitment and differentiation, provided an additional link between the observed BMP-associated response and the broader regulation of BMSC fate(Miraoui and Marie, 2010; Quarto et al., 2015). Previous functional studies demonstrated that TWIST1 silencing enhances osteogenic differentiation through activation of BMP and ERK/FGF signalling and subsequent upregulation of TAZ(Quarto et al., 2015).

In our osteogenic cultures, Si-HAp-LEG-221 significantly reduced *TWIST1* expression, a directionally consistent pattern suggesting attenuation of a transcriptional restraint on osteogenic signalling. However, this response was not accompanied by enhanced mineralization or altered accumulation of osteogenesis-associated proteins, indicating that it remained confined primarily to the regulatory level. Notably, the biomaterial-induced *TWIST1* expression pattern under chondrogenic and adipogenic conditions partially paralleled that of *DANCR*, suggesting coordinated modulation of regulators associated with mesenchymal cell-state control following the Si-HAp platforms doping and co-doping with the Li⁺, Eu³⁺, and Gd³⁺ ions.

It is noteworthy that Si-HAp-LEG-222, which contains a higher proportion of Gd³⁺ ions, consistently elicited the most pronounced molecular response in terms of activation of pro-regenerative transcripts of tissue-specific differentiation. Previous studies have shown that Gd^3+^ ion-doping biomaterials, particularly when combined with other rare-earth elements such as Eu³⁺ ions or HAp-based systems doped or co-doped with rare-earth ions, can enhance osteogenic activity and modulate BMP-related signalling (Zhu et al., 2019; Pan et al., 2025; Moghanian et al., 2026). In this context, the more pronounced molecular response elicited by Si-HAp-LEG-222 may be associated with its higher Gd³⁺ ion content and with composition-dependent interactions between the dopant ions.

Substitution of the Li⁺, Eu³⁺, and Gd³⁺ ions into the Si-HAp biomaterials also affected the post-transcriptional regulatory layer, as reflected by altered *miR-106a* and *miR-30a* expression. Both miRNAs have been implicated in the control of MSC lineage commitment (Tian et al., 2016; Wu et al., 2023). Of particular relevance, *miR-106a* has been shown to regulate osteogenic–adipogenic commitment through BMP2-dependent mechanisms (Li et al., 2013), whereas *miR-30a* has been linked to both *RUNX2*-associated osteogenic regulation and *DLL4/Notch*-dependent chondrogenesis (Tian et al., 2016). Indeed, under osteogenic conditions, the Si-HAp-LEG-222-induced reduction in both miRNAs coincided with a trend toward reduced *RUNX2* transcript levels, whereas their significant upregulation during chondrogenesis revealed increased *RUNX2* mRNA expression. Under adipogenic conditions, cells cultured on the Li⁺, Eu³⁺, and Gd³⁺ ions co-doped biomaterials had reduced *miR-106a* expression, yet this change was not accompanied by altered *RUNX2* transcription. Given that miRNAs are dynamic and sensitive regulators capable of simultaneously influencing multiple target transcripts (Sikora et al., 2020), these findings and molecular signature point to a broader reorganisation of post-transcriptional control, in which the same miRNA may engage distinct molecular axes depending on the differentiation program. Importantly, however, the observed post-transcriptional shifts were not mirrored by corresponding changes in protein abundance, either for lineage-associated regulators or for structural and matrix-related proteins such as type I and type II collagen or in OPG, a regulator relevant to both cartilage homeostasis and bone remodelling (Elango et al., 2018; Wu et al., 2024). These proteins represent two interconnected dimensions of tissue regeneration: type I and type II collagens establish lineage-specific matrix architecture, whereas OPG functionally is associated with tissue remodelling. For example, in nHAp-based systems, coordinated upregulation of COL1, BMP2, RUNX2, and OPG has been associated with enhanced osteogenesis, while OPG silencing attenuated this response (Xu et al., 2026). Viewed in this context, the unchanged abundance of these proteins in osteogenic and chondrogenic BMSC cultures modulated by the presence of Si-HAp biomaterial supports the view that the Li⁺, Eu³⁺, and Gd³⁺ ions co-doping did not disrupt the balance between matrix formation and remodelling.

A similar stability was observed during adipogenesis, where adiponectin and leptin- key adipokines associated with the metabolic and endocrine maturation of BMSC-derived adipocytes - remained unaffected by either the pure Si-HAp or its ions co-doped variants, indicating that biomaterials co-doped with the Li⁺, Eu³⁺, and Gd³⁺ ions did not substantially alter the acquisition of the adipocyte-associated protein phenotype. This finding is relevant beyond the assessment of adipocyte maturation, as bone marrow adipocytes form an endocrine component of the marrow niche, and adiponectin and leptin participate in the metabolic communication between adipose and skeletal tissues (Tencerova and Kassem, 2016; Burkhardt et al., 2023; Zhou et al., 2026).

To summarize, the preserved multilineage competence and stable effector phenotype of BMSCs exposed to the Li⁺, Eu³⁺, and Gd³⁺ ions co-doped Si-HAp biomaterials point to a biologically permissive platform that can accommodate local regenerative signals while providing additional multifunctional and imaging-related properties. From a regenerative perspective, the value of such multifunctional biomaterial lies not only in its ability to induce a strong cellular response, but also in its capacity to remain compatible with the changing demands of the tissue niche.

A material that rigidly enforces a single cellular fate may therefore be less adaptable to the complexity of tissue regeneration. Accordingly, the ions co-doped Si-HAp biomaterials may be viewed not as a strongly lineage-directing material, but as a versatile, niche-compatible platform capable of supporting context-dependent tissue repair. Substitutions of the Li⁺, Eu³⁺, and Gd³⁺ ions to the Si-HAp biomaterials may offer a versatile platform for regenerative medicine and theranostics by supporting cellular responses to the local tissue environment without substantially altering the intrinsic fate of differentiating cells.

## 5 Conclusions

The present study shows that substitutions of Li⁺, Eu³⁺, and Gd³⁺ ions into the Si-HAp biomaterial structure do not diminish the material’s intrinsic capacity to support BMSC regenerative activity. Both Si-HAp-LEG-221 and Si-HAp-LEG-222 remained compatible with osteogenic, chondrogenic, and adipogenic development, while leaving lineage-associated protein profiles and mature phenotypic outcomes unaffected. At the molecular level, however, the ion substitution in the Si-HAp-LEG-222 modified selected transcriptional and post-transcriptional networks involved in tissue-specific commitment, highlighting the potential of this platform to subtly modulate tissue-specific regenerative responses without overriding intrinsic lineage programs. Thus, the Si-HAp co-doped with lithium and rare-earth ions broadens its functionality by adding imaging-related properties without compromising its underlying pro-regenerative character.

## Supporting information

Graphical Abstract

## 7 Author Contributions

Conceptualization, A.Ś., RJ.W., N.C. and K.M.; methodology, N.C., K.M., A.Ś. and J.S.; Validation, A.Ś., RJ.W., N.C., K.M., and A.P.; formal analysis, A.P., A.Ś., K.M.; investigation, A.Ś., K.M., J.S.; resources, A.Ś., RJ.W.; data curation, K.M., A.Ś., A.P.; writing - original draft preparation, A.P., A.Ś. and K.M.; writing - review and editing, A.Ś., RJ.W., K.M., A.P, N.C..; visualization, K.M., A.P.; supervision, A.Ś. and RJ.W; project administration, A.Ś., RJ.W.; funding acquisition, RJ.W. and A.Ś.

All authors have read and agreed to the published version of the manuscript.

## 9 Data Availability Statement

All the data that support the findings of this study are available from the corresponding author upon reasonable request.

## 11 Supplementary Material

**Table S1.**
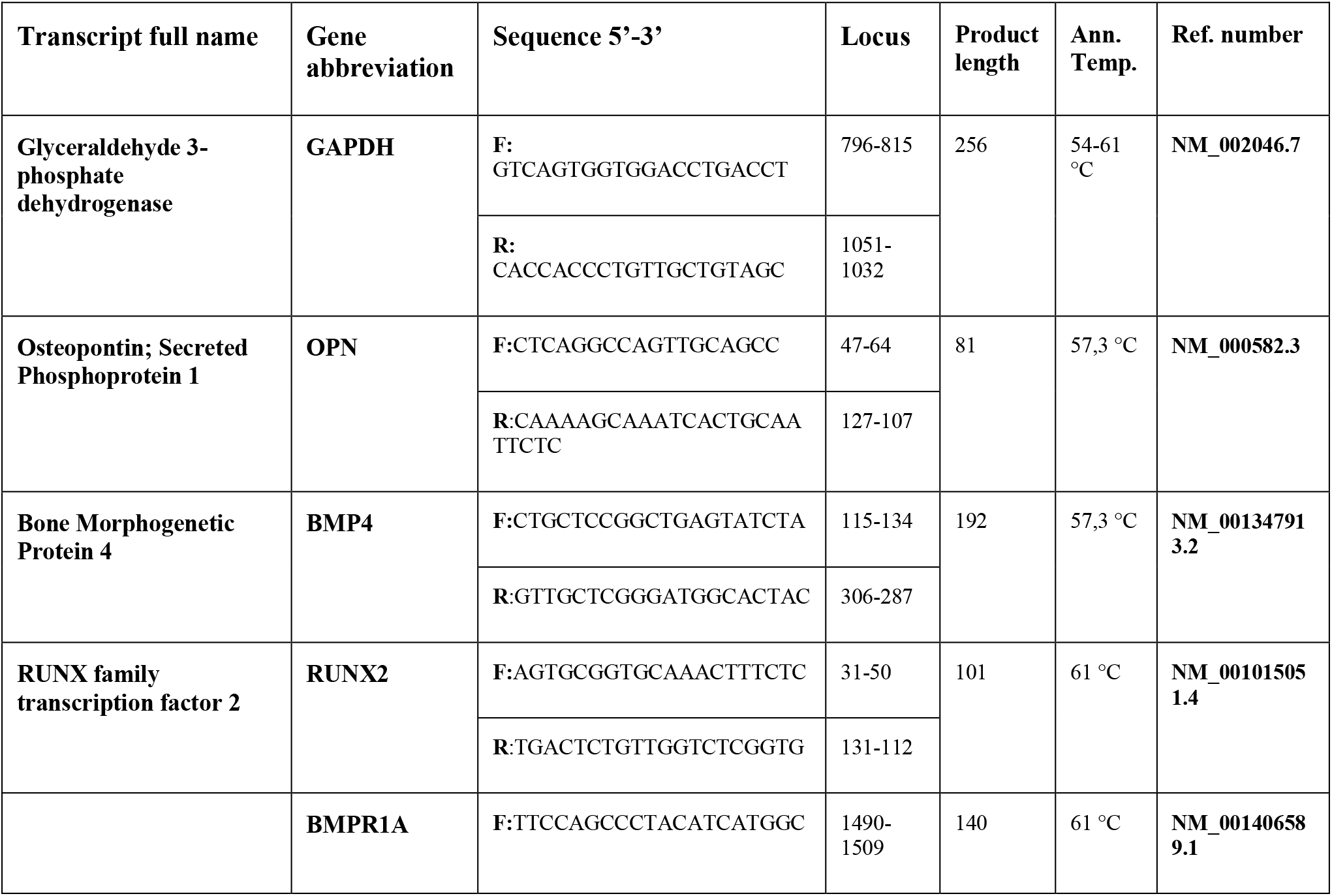
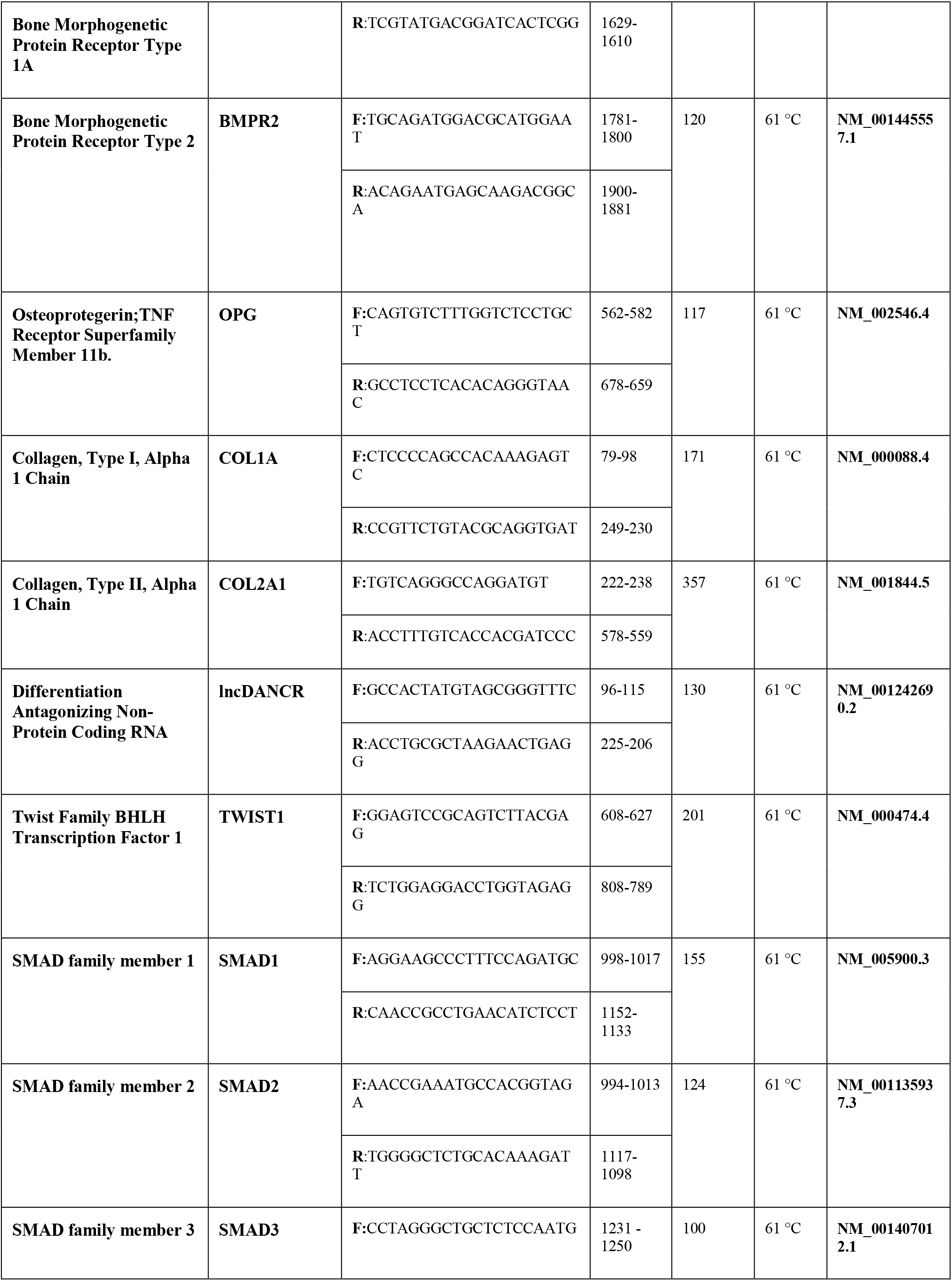
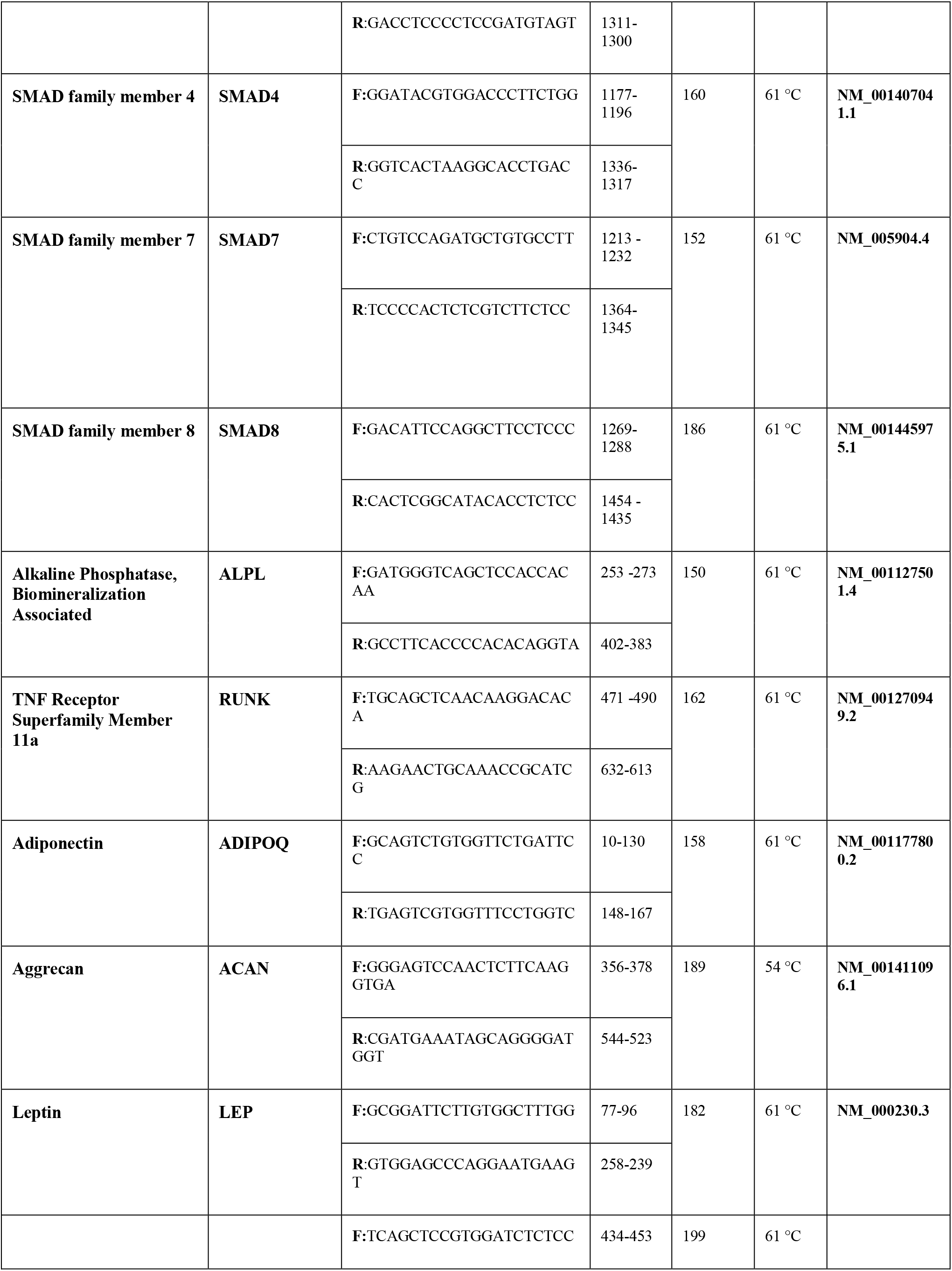
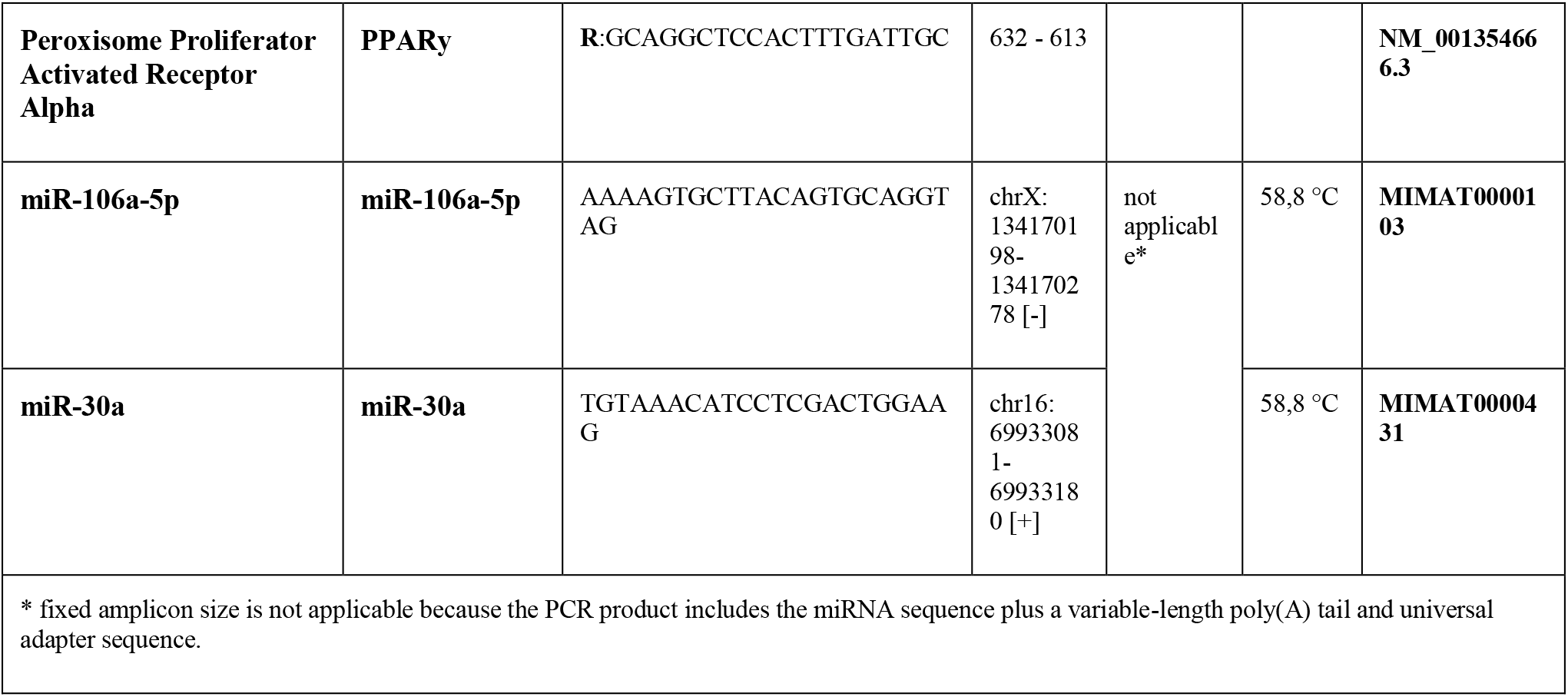
Detailed characteristics of oligonucleotide primers used for RT-qPCR analysis.

**Table S2.** List of antibodies used for Western blot analysis, including the applied working dilutions.

| **Protein name** | **Abbreviation** | **Working dilution** | **Product number** | **Company** |
| --- | --- | --- | --- | --- |
| **Alpha 1 type I collagen** | **COL1** | 1/500 | AF7001 | Affinity Biosciences |
| **Alpha 1 type II collagen** | **COL2** | 1/500 | AF0135 | Affinity Biosciences |
| **Aggrecan 1** | **ACAN** | 1/1000 | DF7561 | Affinity Biosciences |
| **Core binding factor subunit alpha 1** | **RUNX2** | 1/500 | AF5186 | Affinity Biosciences |
| **Osteoprotegerin** | **OPG** | 1/500 | DF6824 | Affinity Biosciences |
| **Osteopontin** | **OPN** | 1/500 | AF0227 | Affinity Biosciences |
| **Bone morphogenetic protein 2** | **BMP2** | 1/500 | AF5163 | Affinity<br>Biosciences |
| **Adipocyte complement related protein** | **Adiponectin** | 1/500 | DF7000 | Affinity<br>Biosciences |
| **Leptin** | **Leptin** | 1/1000 | DF8583 | Affinity<br>Biosciences |
| **Beta Actin** | **ACTB** | 1/5000 | AF7018 | Affinity<br>Biosciences |

