## Supplementary figures and images for "Multifunctional Rare-Earth-Ion-Doped Si-HAp Platforms Modulate Human BMSC Lineage-Associated Molecular Responses Without Enhancing Terminal Differentiation"

### Graphical Abstract

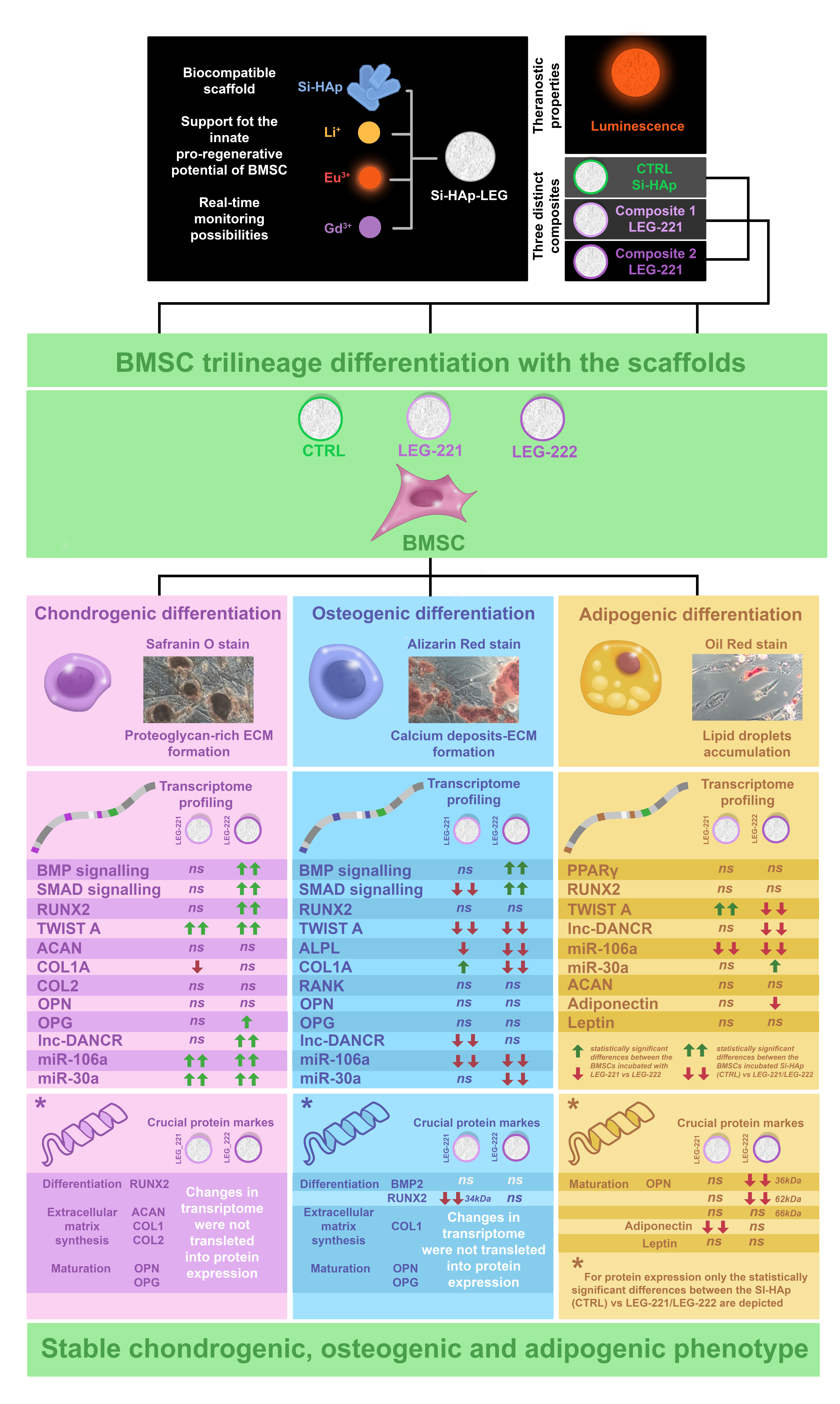
